# eXNVerify: coverage analysis for long and short-read sequencing data in clinical context

**DOI:** 10.1101/2021.12.16.473078

**Authors:** Sebastian Porebski, Tomasz Stokowy

## Abstract

Accurate identification of genetic variants to a large extent is based on type of experimental technology, quality of the material and coverage of obtained sequencing data. Our motivation was to create a tool that will evaluate genome coverage and accelerate the introduction of long-read sequencing to medical diagnostics and clinical practice. Here we present eXNVerify: a tool for inspection of clinical data in the context of pathogenic variants. The tool calculates Clinical Depth Coverage – a measure of coverage which we introduce to evaluate loci with pathogenic germline and somatic variants reported in ClinVar. The tool additionally provides visualization options for user-defined genes of interest. Finally, we present an examples of BRCA1, TP53, CFTR application and results of a test conducted in the Extensive Sequence Dataset of Gold-Standard Samples for Benchmarking and Development. eXNVerify is available at https://github.com/porebskis/eXNVerify and can be directly pulled from the DockerHub repository: docker pull porebskis/exnverify:1.0.

## Introduction

Accurate identification of clinically relevant genomic variants strictly depends on sequencing coverage of sequencing data. Long-read sequencing covers a higher percentage of the human genome than short-read sequencing and results in more stable coverage [1]. Consequently, single nucleotide, indel [2], and structural variants are detected more accurately. Recent benchmarks evaluate the accuracy, precision, and recall of variant calling in long-read genome data [4], however, the introduction of new findings in the clinical and diagnostic setting requires more time. To accelerate the development of clinical genomics we present eXNVerify named from “exon and single nucleotide variant (SNV) verification”), a standalone tool that evaluates and visualizes genome coverage in a clinical context. While the available software approaches can analyze the sequencing data, none of them focuses on evaluating SNV coverage in the context of diagnostic procedures. Our goal was to fill this gap is filled by eXNVerify. The comprehensive quality control of medically relevant genes can be now adjusted to the diagnostic procedure. Moreover, the tool helps to verify the sequencing sample in terms of coverage of selected genes or to evaluate the overall genome/exome in terms of variant coverage.

## Methods

eXNVerify tool is designed to run clinically relevant coverage analysis for SR and LR data, providing integration with the ClinVar database. The software is designed as two standalone procedures: geneCoverage and snvScore. The primary input file for both procedures is the coverage record (BED format) obtained from processing BAM files. To create an input file, the user can use dependencies such as bedtools [5], mosdepth [6], or samtools [7] (see GitHub documentation: https://github.com/porebskis/eXNVerify).). The first procedure is geneCoverage. According to the location of exons of the selected gene, it presents coverage in a graphical form (coverages of exons are light blue fragments in Fig 1) with the location of pathogenic SNVs. Germline and somatic variants are shown as red and dark blue dots, respectively. Moreover, geneCoverage counts the coverage of these SNVs and summarizes the results in a tabular form. The geneCoverage script, in addition to the exon list, pathogenic germline, and pathogenic somatic SNV list, also takes the names of the genes and the coverage threshold as a parameter. The latter is set to evaluate the sample if the gene-related variants are sufficiently covered. Thus, geneCoverage reports the percentage of SNV covered for a given gene and includes insufficient coverage in the generated figure. That is, specific exons that are poorly covered (which may contain key variants) are highlighted in red (see Fig 1, 2 and 3).

**Fig 1.**
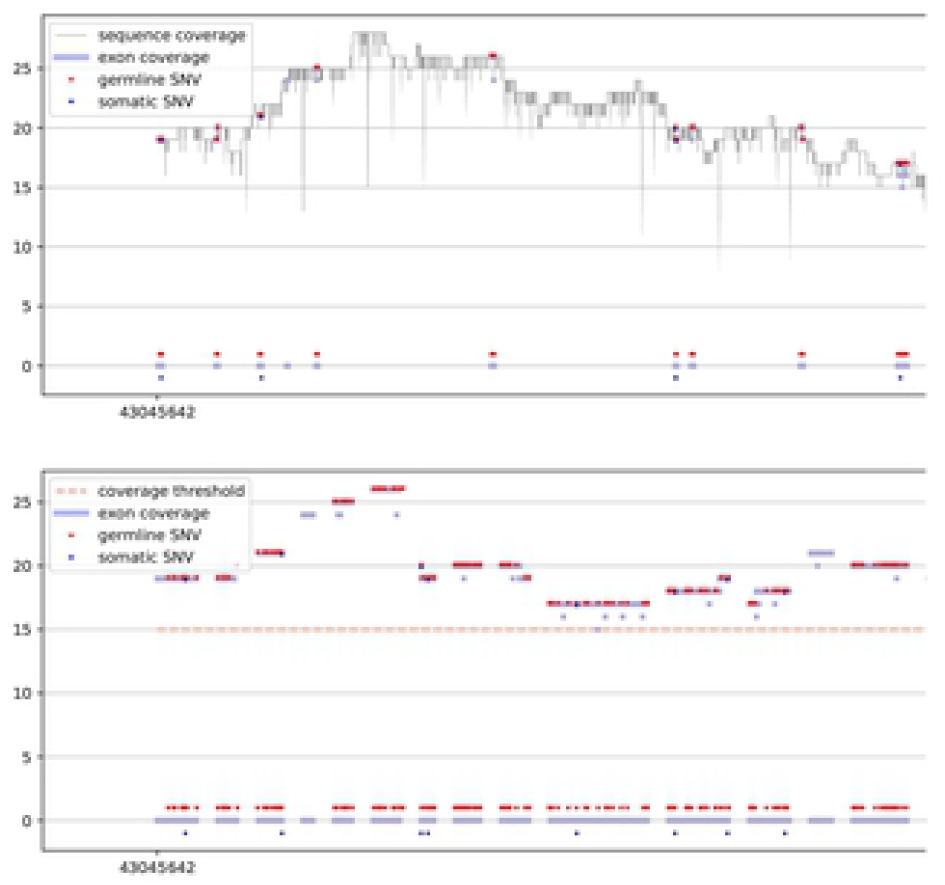
BRCA1 coverage for sample PacBio Long Read. The upper panel demonstrates the distribution of coverage in the region of the gene (exons and introns). The lower panel depicts coverage in exons. X-axis is a genomic locus, specified by the user. Dots highlight positions where pathogenic germline (red) and somatic (dark blue) ClinVar variants are located. If coverage of exons is lower than the threshold specified by the user, they are highlighted by the software in red color.

**Fig 2.**
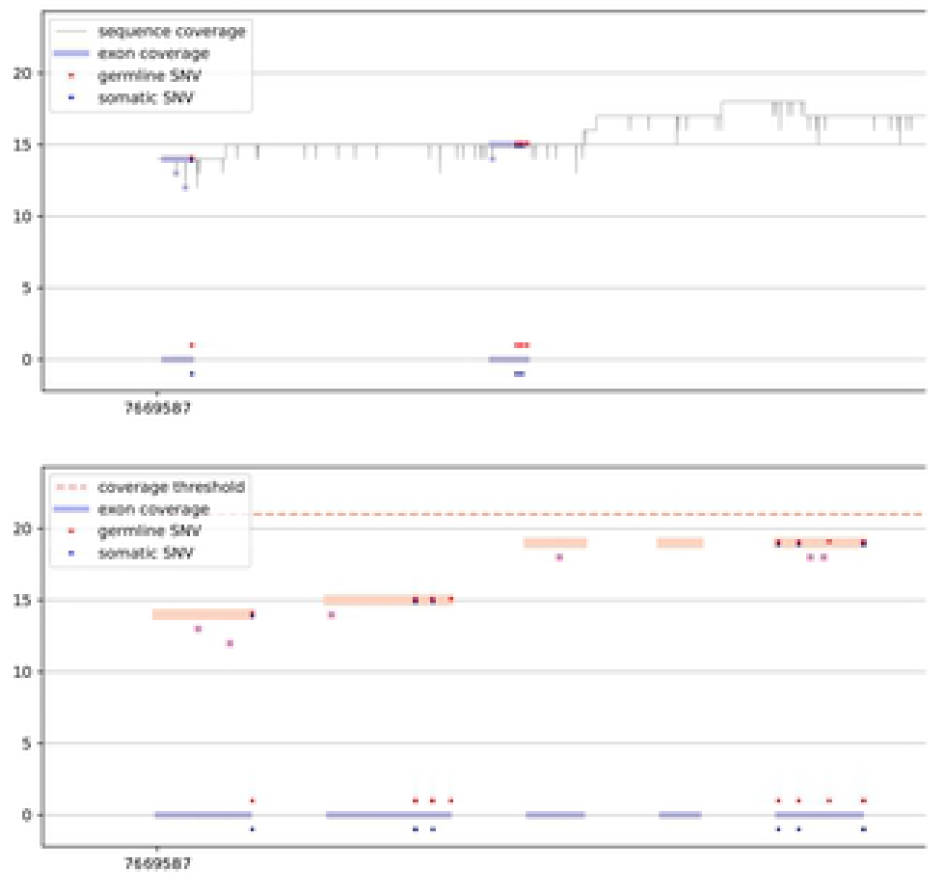
TP53 coverage for PacBio Long Read sample. Coverage of more than 40% of the gene did not reach expected 20x coverage. In such case diagnostic lab should consider optimization of the sequencing protocol, especially in exons 1, 2, 5 and 6, which include germline and somatic pathogenic variants.

**Fig 3.**
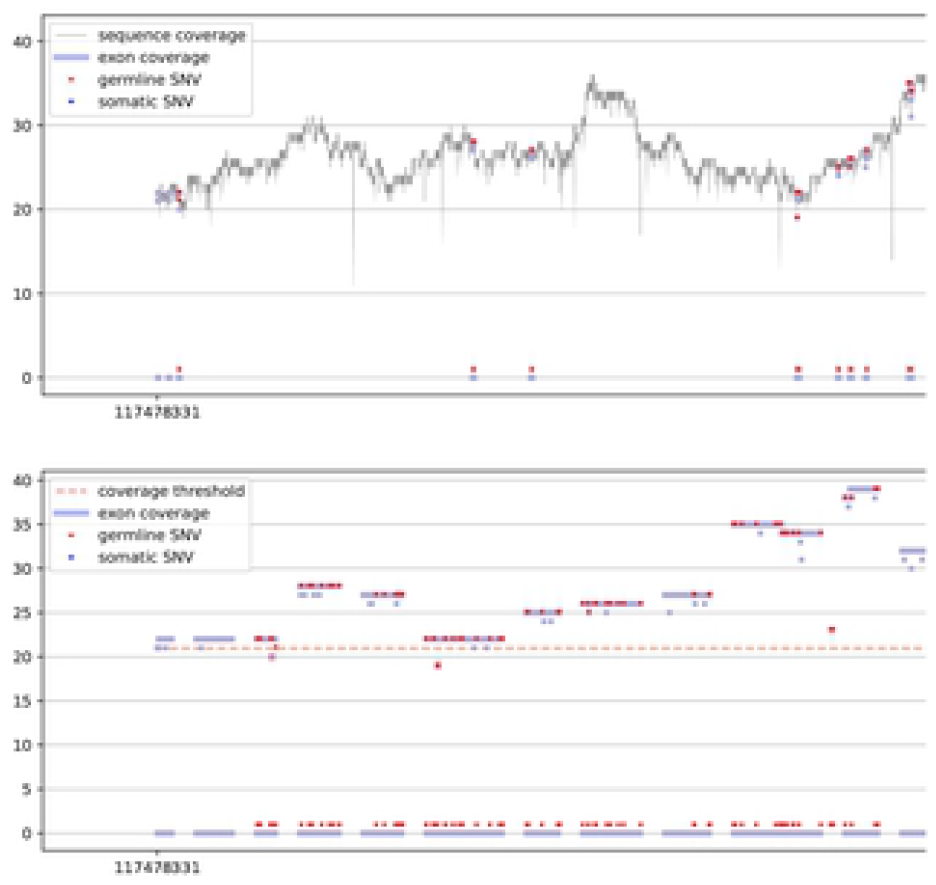
CFTR coverage for PacBio Long Read sample. eXNVerify scales visualization to illustrate correctly genes with large number of exons. In this case, CFTR gene, responsible for development of cystic fibrosis consist of 27 exomes.

This design helps to evaluate the reliability of the data before and after specific variant calling. Importantly, it is possible to prepare their own reference files with desired exon regions and SNV positions by following the examples provided in referenced GitHub repository.

An additional element of eXNVerify that focuses on the overall evaluation of sequence coverage is snvScore. It is used to check the coverage of all SNVs downloaded from the ClinVar database. The snvScore script checks variant coverage by all chromosomes and provides basic statistics. Finally, snvScore calculates a proposed measure of variant coverage, called Clinical Depth Coverage, calculated as:

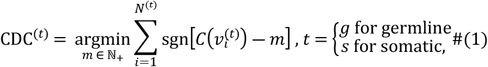

where 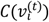 is the coverage of *i*-th SNV and *N*^(*t*)^is the number of all referenced variants. Germline and somatic variants are analyzed separately, hence *t* equals g or s for germline and somatic, respectively. We created examples for genes highly relevant for medical genetics and cancer genomics (see Supplementary Files 1 and 2).

### Requirements

Both scripts are included in Docker container eXNVerify, ready for pulling from DockerHub repository. Furthermore, both scripts can be executed in the user system if Python 3.8 with numpy, pandas and matplotlib libraries are established. Execution of Docker container is widely described in GitHub documentation and Supplementary Data for this work. Below, general requirements of input files and processing pipeline are included.

### User input

Both scripts, geneCoverage, and snvScore take a similar set of input files: sample BED file (per-base analysis result from BAM file), a reference list of exons (BED format, only for geneCoverage), two ClinVar-downloaded tables with pathogenic germline and somatic SNVs (text table). For both scripts, a user may choose a coverage threshold. For geneCoverage, an additional required parameter is the list of genes (at least one gene name).

### Implementation in detail

Sample BED file contains exclusive fragments of a gene sequence. Each fragment is related to one value of coverage. An exome reference list is also a BED format file, but it is different since it contains the location of all exons. It means that each row usually expresses a large fragment of sequence. Hence one fragment may be covered in a different number of reads. The task was to locate all fragments in the sample BED file that are in one exon fragment from reference BED. Coverages in one exon fragment are extracted. Next, if SNV location is available, it is possible to extract coverage information in formerly extracted exon coverage information. Coverage of SNVs is also presented graphically, but SNV coverage information is aggregated and summarized in table form in a report file. These operations are the core of the geneCoverage procedure.

The second procedure, snvScore explores the whole genome/exome and extracts all referenced SNVs coverage. Hence, snvScore requires a sample BED file, two ClinVar SNV tables, but exome reference and gene name are not necessary. Sample BED may be a large file hence snvScore iteratively load one-chromosome fragments, extracts, and aggregates information about SNV coverage. When finished, it reports CDC (1) for the whole sample with via-chromosome table of germline and somatic pathogenic SNV coverage statistics.

## Results

eXNVerify is a new tool created to evaluate and visualize gene coverage in a clinical context. The tool consists of two methods implemented in Python: geneCoverage and snvScore. The first tool, geneCoverage looks for a gene (or multiple genes) of interest and evaluates it, integrating the coverage with ClinVar pathogenic variant information. It demonstrates exons in a gene of interest, highlighting positions of pathogenic variants in the ClinVar database (Fig 1, BRCA1 gene). The tool includes both germline pathogenic and somatic pathogenic single nucleotide variants. The tool is flexible and suitable for both oncology project (Fig 2, TP53 gene) and rare disease projects (Fig 3, CFTR gene). Processing the samples with the pandas and numpy libraries as well as visualization of the results with the matplotlib library is enough to provide intuitive support for the diagnostician. Additionally, geneCoverage indicates positions in which desired coverage has not been achieved and therefore variant analysis may lead to false negative/positive calls. The tool is suitable for LR and SR data providing novel insights and analysis options for all technologies used currently in clinical laboratories. To address the spectrum of technology-dependent coverage differences we present results for LR whole genome, SR whole genome and SR exome in Supplementary Fig 1 A, B, and C, respectively. The second method, snvScore calculates coverage statistics for pathogenic variants, allowing the user to estimate the percentage of all SNVs that are covered above the defined threshold. Table 1 summarizes the essentials of its execution for test samples (Supplementary File 3).

**Table 1.**
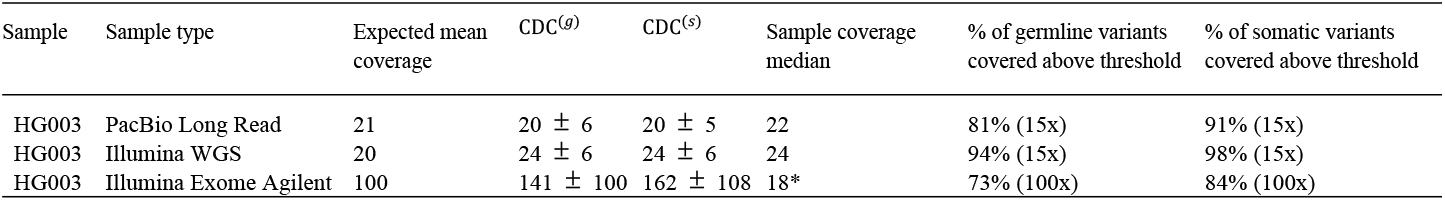
Results obtained for test samples.

The tool can be used to inspect structural variants observed in the sample, especially deletions and copy number changes. This approach can be helpful in a manual verification of structural variants, which is still a recommended practice in medical genetics [8]. Finally, eXNVerify gives insight into the sample’s usefulness in a hypothesis-free analysis of pathogenic variants. The proposed Clinical Depth Coverage measure provides a percentage of variants covered above the desired threshold in a specified case (Supplementary Table 1 and Supplementary File 3). This measure is useful for everyday laboratory practice to maintain and maximize the quality of experiments. Results of such analysis are provided in Supplementary Table 1, which indicates % of germline and somatic pathogenic variants specified above the desired threshold. It can be also observed that pathogenic variant coverage differs from median coverage and mean coverage of the sample. For the HG003 PacBio Long Read sample, CDC equals 20x, while global median coverage is 22x. A user of the software can also see that 81% of germline pathogenic variants were covered at least 15x. We conclude that CDC measures and the percentage of variants covered above the threshold are useful for medical genetics and cancer diagnostics. In summary, our new tool introduces new, easily applicable options for medical genome analysis.

## Supporting information

**S1 Fig BRCA1 coverage for samples: A – PacBio Long Read, B – Illumina WGS, C – Illumina Exome Agilent**. Detail description as for Fig 1.

**S1 File Supplementary Data**

**S2 File geneCoverage report for HG003 PacBio LR, Illumina WGS, Illumina Exome Agilent**

**S3 File snvScore report for HG003 PacBio LR, Illumina WGS, Illumina Exome Agilent**

### Availability

Ready-to-go Docker container with eXNVerify can be pulled directly from DockerHub using the following command: docker pull porebskis/exnverify:1.0. Detailed documentation of the project, exemplar execution results, and source code are available at: https://github.com/porebskis/eXNVerify.

## Acknowledgements

We would like to acknowledge SnotraBio for sharing computational resources which were used to develop the study. We are thankful for constructive insights from employees of the Medical Genetics Department, Haukeland University Hospital, Bergen Norway, especially from Aashish Srivastava, Rita Holdhus and Sigrid Erdal.

